# Epithelial organoid supports resident memory CD8 T cell differentiation

**DOI:** 10.1101/2023.12.01.569395

**Authors:** Max R. Ulibarri, Ying Lin, Julian R. Ramprashad, Geongoo Han, Mohammad H. Hasan, Farha J. Mithila, Chaoyu Ma, Smita Gopinath, Nu Zhang, J. Justin Milner, Lalit K. Beura

## Abstract

Resident Memory T cells (TRM) play a vital role in regional immune defense in barrier organs. Although laboratory rodents have been extensively used to study fundamental TRM biology, poor isolation efficiency, sampling bias and low cell survival rates have limited our ability to conduct TRM-focused high-throughput assays. Here, we engineered a murine vaginal epithelial organoid (VEO)-CD8 T cell co-culture system that supports CD8 TRM differentiation *in vitro*. The three-dimensional VEOs established from murine adult stem cells resembled stratified squamous vaginal epithelium and induced gradual differentiation of activated CD8 T cells into epithelial TRM. These *in vitro* generated TRM were phenotypically and transcriptionally similar to *in vivo* TRM, and key tissue residency features were reinforced with a second cognate-antigen exposure during co-culture. TRM differentiation was not affected even when VEOs and CD8 T cells were separated by a semipermeable barrier, indicating soluble factors’ involvement. Pharmacological and genetic approaches showed that TGF-β signaling played a crucial role in their differentiation. We found that the VEOs in our model remained susceptible to viral infections and the CD8 T cells were amenable to genetic manipulation; both of which will allow detailed interrogation of antiviral CD8 T cell biology in a reductionist setting. In summary, we established a robust model which captures bonafide TRM differentiation that is scalable, open to iterative sampling, and can be subjected to high throughput assays that will rapidly add to our understanding of TRM.

## Introduction

Memory CD8 T cells play a crucial role in coordinating the immune response against intracellular infections and malignancies. Their duties, however, are compartmentalized, with distinct subsets of memory CD8 T cells performing surveillance responsibilities depending on their anatomic location. Specifically, circulating memory CD8 T cells, which encompass both central memory (TCM) and effector memory CD8 T cells (TEM), continuously patrol the bloodstream and secondary lymphoid organs (SLOs) such as the spleen and lymph nodes as well as non-lymphoid tissues (NLTs) ^1–3^. Resident memory CD8 T cells (TRM), by contrast, are stationed within specific tissues and rarely recirculate through blood or lymphatics. The frontline placement of TRM positions them at the first line of defense against invading pathogens. Upon contact with infected Antigen Presenting Cells, TRM cells promptly release a milieu of cytokines and chemokines and exhibit cytotoxic capacity. This multifaceted response serves to curtail pathogen replication, alert the immune system, and recruit other immune cells to the site of infection. Consequently, the presence of TRM is correlated with expedited pathogen control in a number of barrier tissues ^4,5^.

Mucosal barrier tissues including the intestine, lung and female reproductive tract (FRT) are frequently targeted by pathogens. Localizing abundant quantities of antiviral CD8 TRM in these tissues is associated with rapid protective benefit in infection ^6–10^. Accordingly, positioning a robust TRM population in barrier tissues that is maintained long-term is a crucial vaccination goal. This requires an in-depth understanding of the signals that mediate differentiation of naive CD8 T cells to TRM While the identity of certain core transcription factors (e.g. Hobit, Blimp-1, Runx3 and KLF2) and surface molecules (e.g. CD103, CD69, CD49a) have been discovered, our understanding of the TRM differentiation process is far from complete ^11–13.TRM^ development is complex and involves multiple anatomical niches including initial effector differentiation in SLOs, trafficking via blood, and final TRM formation at the tissue of residence under the influence of the local microenvironment. The contributions of the local tissue-specific signals in dictating TRM fate is an intense area of research as the information could be used to modulate TRM density in an organ-restricted manner. Many of these studies employ gene-specific knockout mice and transgenic CD8 T cells to elucidate mechanistic insights into the signaling mechanism that induces TRM. However, a major issue remains in distinguishing the roles of specific genes in the initial CD8 T cell effector differentiation process, which occurs in SLOs, from their contributions to the subsequent differentiation process that transpires within the respective non-lymphoid barrier tissues, once the T cells have homed there. The utility of tissue-specific Cre-driver lines, which can be temporally induced, is constrained by their limited availability and susceptibility to spurious or leaky induction. Furthermore, these *in vivo* animal studies are not well-suited for high-throughput assays and are constrained in their capacity for invasive experimental manipulations. Addressing these limitations, organoid models have emerged as a reductionist surrogate system that overcomes the shortcomings of *in vivo* models while retaining the three-dimensional architecture of target tissues ^14–16^.

Epithelial organoids can be derived from induced pluripotent stem cells or adult epithelial stem cells. They are phenotypically stable through successive passages, which makes them an efficacious alternative to *in vivo* assays. Other components of the native tissue can also be incorporated into these epithelial organoids including immune cells, mesenchymal cells, and a microbiome to develop more faithful models that recapitulate relevant *in vivo* interactions ^17–19^. Enteric and lung organoids have been well-established and currently offer tremendous prospect for fundamental biologic discovery as well as personalized medicine. In comparison, organs with type-II mucosa have been less investigated. Here we exploited a recently established model of vaginal epithelial organoids (VEO) ^20^ to dissect the localized interactions between T cells and the vaginal epithelium and to study TRM differentiation. By co-culturing activated CD8 T cells with VEOs, we successfully induced CD8 TRM differentiation. Subsequent analysis of the transcriptome and phenotype of the CD8 T cells showed robust alignment of the *in vitro* generated TRM with bonafide *in vivo* CD8 TRM cells. We further ascertained that the TRM phenotype is contingent on TGF-β signaling and can be repressed by the chemical inhibition of TGF-β activation/signaling. This reductionist model system enables in-depth exploration of the intricate interplay between T cells and the vaginal epithelium, providing valuable insights into the local differentiation of TRM within the FRT.

## Results

### Establishment of vaginal epithelial organoid-CD8 T cell co-culture system

In this study, we employed a vaginal epithelial organoid (VEO) generation system, as previously outlined by Ali et al. in 2020^21^, to cultivate VEOs. Vaginal tissue was collected from female C57BL/6 mice (>8 weeks), and the epithelium was separated from stroma using a combination of enzymatic and physical techniques (Fig. 1A). Single-cell suspensions of epithelial cells were embedded in basement membrane extract (Matrigel) and cultured in a growth medium designed for the maintenance and proliferation of epithelial stem cells. Individual stem cells gave rise to structures with multiple differentiated layers, closely resembling previously described vaginal organoids^20^. These VEOs were successfully maintained for at least 21 days through supplementation of fresh medium during which they steadily grew in size (Fig.1B). Notably, in differential interference contrast (DIC) images, the organoids exhibited a distinct darker core attributed to mucinous secretions and a lighter external boundary composed of a basal layer of epithelial cells (Fig.1B). We also measured relative transcript levels of various genes associated with different layers of vaginal epithelium at different times post-culture ^20^. Transcripts associated with stem cells (*Axin2*) and proliferation (*Birc5, Ki67*) were more abundant at earlier times (day 5 post-culture), whereas genes associated with luminal keratinocytes (*Sprr1a*) and cornified cells (*Krt1*) increased at later times (Fig.1C) ^20,22^.

**Figure 1:**
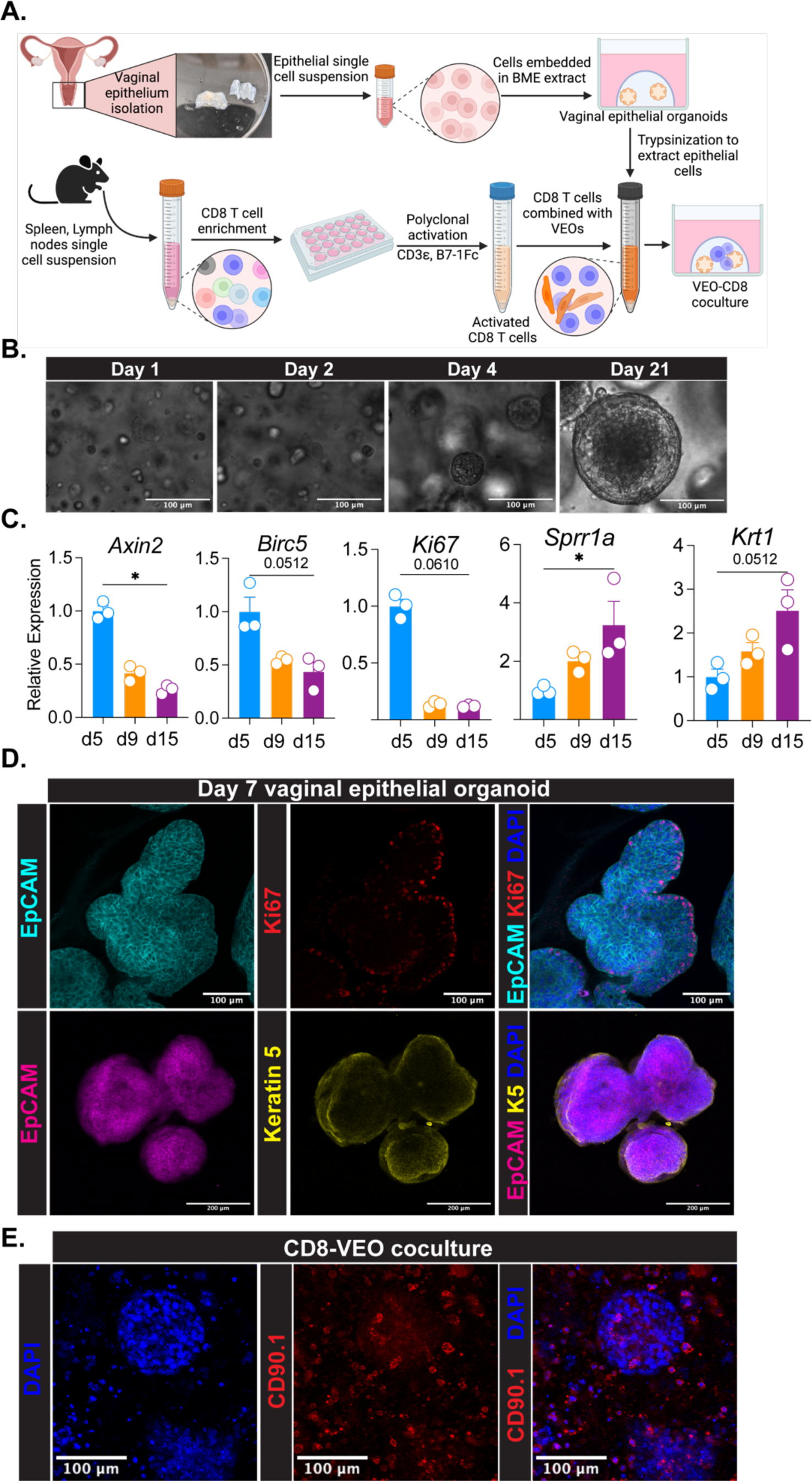
Establishment of vaginal epithelial organoid (VEO) and co-culturing with CD8 T lymphocytes. **A.** Schematics describing the isolation of vaginal epithelial cells from C57BL/6 mouse and differentiation of epithelial organoid using growth factors and chemicals. Naive CD8 T cells from TCR transgenic mice were enriched and activated *in vitro* using CD3 and B7.1 Fc. Activated CD8 T cells were co-cultured with VEOs to enable CD8 T cells’ differentiation to TRM. **B.** Representative differential interference contrast (DIC) microscopy of VEOs at day 1, 2, 4 and 21 post subculture showing growth. Scale bar=100µm. C. Relative RNA level of indicated genes detected by quantitative PCR at different days post-subculture showing differential levels of distinct epithelial populations within the VEOs as they grow. **D.** Representative confocal microscopy images of VEOs showing epithelial identity as well as different layers. Top row-Epcam, cyan; Ki67, red; DAPI, blue; Scale bar=100µm. Bottom row-Epcam, magenta; Keratin 5, yellow; DAPI, blue; Scale bar=200µm. **E**. Activated CD8 T cells stained with a congenic marker CD90.1 (red) were co-cultured with VEOs, and representative confocal microscopy image 7-day post culture is shown. Scale bar=100µm. Schematic in A is made with Biorender. Experiments in B, C, D and E have been repeated at least twice with more than 3 separate wells/condition.

Our histological analysis of the VEOs demonstrated consistent staining with the pan-epithelial cell marker EpCAM. The majority of proliferating cells (Ki67+) were located in the outer layer, (Fig. 1D, top row). Similarly, the basal epithelial cell marker, keratin-5, was predominantly localized to the outer layer of cells within the organoids (Fig. 1D, bottom row). Next, we aimed to introduce CD8 T cells into the VEOs to test whether exposure to VEO-derived cues could facilitate CD8 T cell differentiation into mature tissue-resident memory (TRM) cells. To achieve this, naive CD8 T cells were enriched from secondary lymphoid organs (SLOs) of TCR transgenic mice, which included P14 mice carrying CD8 T cells with specificity for the gp33 epitope of Lymphocytic choriomeningitis virus (LCMV), OT-I mice bearing CD8 T cells against the ovalbumin-derived SIINFEKL epitope, or gBT-I mice containing CD8 T cells with specificity for the SL8 epitope of Herpes simplex virus (HSV). These cells underwent polyclonal activation using a combination of a CD3ε antibody and B7.1 Fc as described before ^23^. The expanded CD8 T cells were then co-cultured with the VEOs, as shown in Fig. 1A, and their presence within the basement membrane extract (BME)/Matrigel was imaged using a congenic marker (CD90.1) through confocal microscopy. CD8 T cells were observed in close proximity to fully developed organoids as well as found scattered around the organoids, as presented in Fig. 1E. These CD8-VEO co-cultures were successfully maintained for a minimum of 16 days with regular media changes, supplemented with IL-2 alone. In summary, we established VEOs that closely resemble previously described organoids and effectively introduced CD8 T cells into the VEO environment for further investigation.

### CD8 T cells acquire an epithelial TRM phenotype upon co-culture with VEOs

Following the successful maintenance of CD8 T cells with VEOs, we performed phenotypic characterizations of these co-cultured CD8 T cells. Expression of various CD8 T cell-specific markers were assessed from dissociated VEO-CD8 co-cultures via flow cytometry. CD8 T cells maintained alone in the absence of VEOs upregulated CD103 but not CD69, and few cells expressed both CD69 and CD103 (Fig.2A, top row). In contrast, a substantial proportion (∼40-65%) of the co-cultured CD8 T cells showed dual expression of CD69 and CD103 (Fig.2A, middle row). This double positive CD8 TRM cell population is normally observed in the epithelial compartment ^24–26^. Importantly, this *in vitro* co-cultured CD8 T cell phenotype resembled that of anti-viral CD8 TRM generated against murine LCMV infection *in vivo* (Fig.2A, bottom row). These CD8 T cells are well-documented as bonafide residents within the vaginal and cervical tissues^27,28^. Beyond CD69 and CD103, the co-cultured CD8 T cells also adopted other phenotypic attributes of TRM, including downregulation of Ly6C and CD62L and upregulation of PD-1 ^24,29^Interaction of TRM with extracellular matrix (ECM) components is important for many aspects of TRM biology ^30–32^. To test if the acquisition of the TRM phenotype was influenced by the presence of Matrigel (ECM), which provides a 3D environment and support for the growth and maintenance of VEOs, we cultured CD8 T cells within Matrigel in the absence of VEOs. Even after 12 days of culture, these CD8 T cells failed to adopt a CD69+CD103+ TRM phenotype, confirming that ECM alone cannot drive the TRM phenotype and VEOs are crucial in driving TRM formation (Supplementary Fig.1A-B). Additionally, co-cultured CD8 T cells exhibited downregulation of transcription factors T-bet and Eomes, aligning with established TRM traits (Supplementary Fig.1C)^33^.

**Figure 2:**
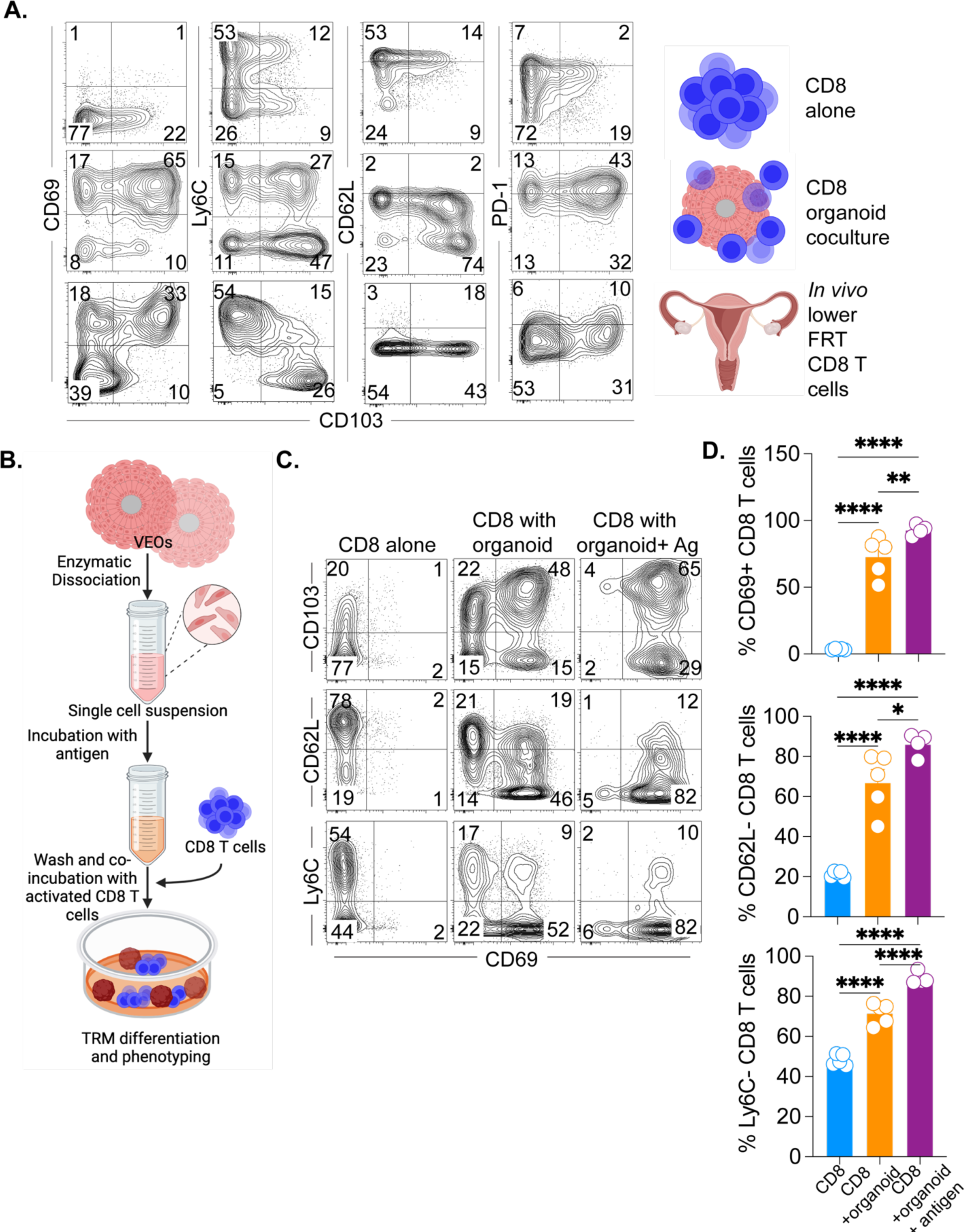
Co-cultured CD8 T cells adopt phenotypic characteristics of TRM. A. CD8 T cells maintained alone (top row) or embedded with the VEOs (middle row) were isolated at day 14 post culture, and representative flow plots depicting expression of various TRM-associated markers are shown. Both rows were gated on live congenic marker (CD45.1 or CD90.1) or CD8β+ T cells. Flow plots in the bottom row are viral antigen-specific memory CD8 T cells isolated from the lower FRT of mice infected with lymphocytic choriomeningitis virus (LCMV) 50 days prior. The plots are gated on live antigen-specific CD8 T cells located in the tissue parenchyma (IV negative). **B.** Schematics describing the protocol used to expose the activated CD8 T cells to cognate antigen again during the co-culture. **C.** Flow cytometry phenotype of CD8 T cells exposed to antigen (gp33 peptide) leading to enhanced acquisition of TRM characteristics. Representative flow plots are shown in C gated on live congenic marker (CD45.1 or CD90.1) or CD8β+ T cells, and percentages are enumerated in **D**. Bars indicate mean ± SEM. Data are representative of three repeats with n=4-6/condition. One way ANOVA with Holm-Sidak multiple comparison test (D). * < 0.05, ** < 0.01, *** <0.001 and ****<0.0001.

To gain insight into the kinetics of acquisition of various TRM markers, we conducted longitudinal phenotyping of co-cultured CD8 T cells, revealing that CD103 upregulation occurs at a faster rate compared to CD69 in vitro (Supplementary Fig.2A-B). We further performed deeper phenotypic characterization of these different CD8 populations generated via co-culture. This revealed that the CD69+ CD103+ CD8 T cells conform to the established true TRM phenotype (CD62L lo, P2rx7 hi, CXCR6 hi) and express CD49a as well as the cytotoxic molecule granzyme-B (Supplementary Fig.2C). However, the CD103 single positive CD8 T cells failed to adopt these TRM phenotypes and rather resembled circulating CD8 T cells expressing higher levels of CD62L.

Previous studies in mouse models have suggested that a local second antigenic encounter in the target tissue can enhance the differentiation of effector CD8 T cells into TRM ^34,35^. In our study, we aimed to replicate this process by exposing the activated CD8 T cells to VEOs presenting cognate antigen. For this, disaggregated epithelial cells from VEOs were incubated with cognate antigenic peptide (gp33 for P14 CD8 T cells and SL8 for gBT-I CD8 T cells) for an hour and were subsequently washed to eliminate any unbound peptides (Fig.2D). Epitope-loaded epithelial cells were incubated with activated CD8 T cells, and together, were embedded in BME to induce VEO formation and TRM differentiation. The antigen-exposed CD8 T cells exhibited a significantly higher percentage of CD69+CD103+ TRM, as depicted in Figure 2E&2F, as early as 8 days in comparison to the non-antigen-exposed CD8 T cells. The expression of other TRM-associated markers was also more pronounced in these cells (Fig. 2E-F). In summary, we have successfully differentiated CD8 TRM cells through VEO-derived signals, and this process was enhanced by a transient second antigen exposure.

**Supplementary Figure 1.**
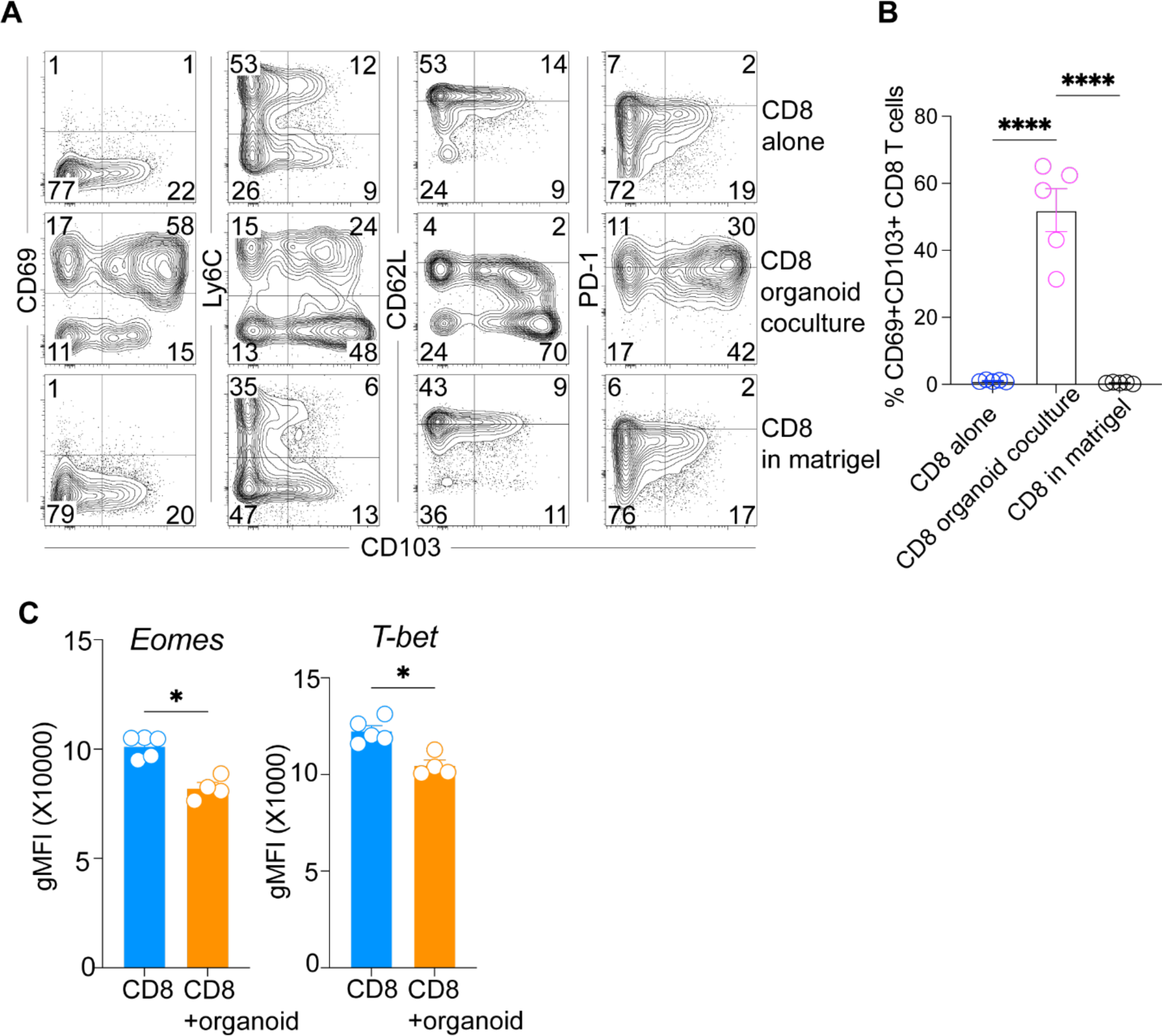
The extracellular matrix microenvironment alone is not capable of supporting TRM differentiation. **A**. CD8 T cells were mixed with VEO-derived epithelial cells and were embedded in Matrigel (middle row) or just embedded alone in BME in the absence of VEOs (bottom row) for 14 days. CD8 T cells maintained as a suspension culture in the absence of VEOs (top row) were included as a control. Representative flow plots depicting expression of various TRM-associated markers are shown. Cells were gated on live congenic marker (CD45.1 or CD90.1) or CD8β+ T cells. **B**. Bar graph comparing the percentage acquisition of TRM phenotype among different conditions. **C**. Geometric mean fluorescence intensity comparison of transcription factors eomesodermin and T-bet between *in vitro* generated TRM and CD8 T cells maintained alone after 14 days of culture. Data are representative of two repeats with n=4-6/condition. Bars indicate mean ± SEM. One way ANOVA with Holm-Sidak multiple comparison test (B). Student t-test (C). * < 0.05, ****<0.0001

**Supplementary Figure 2.**
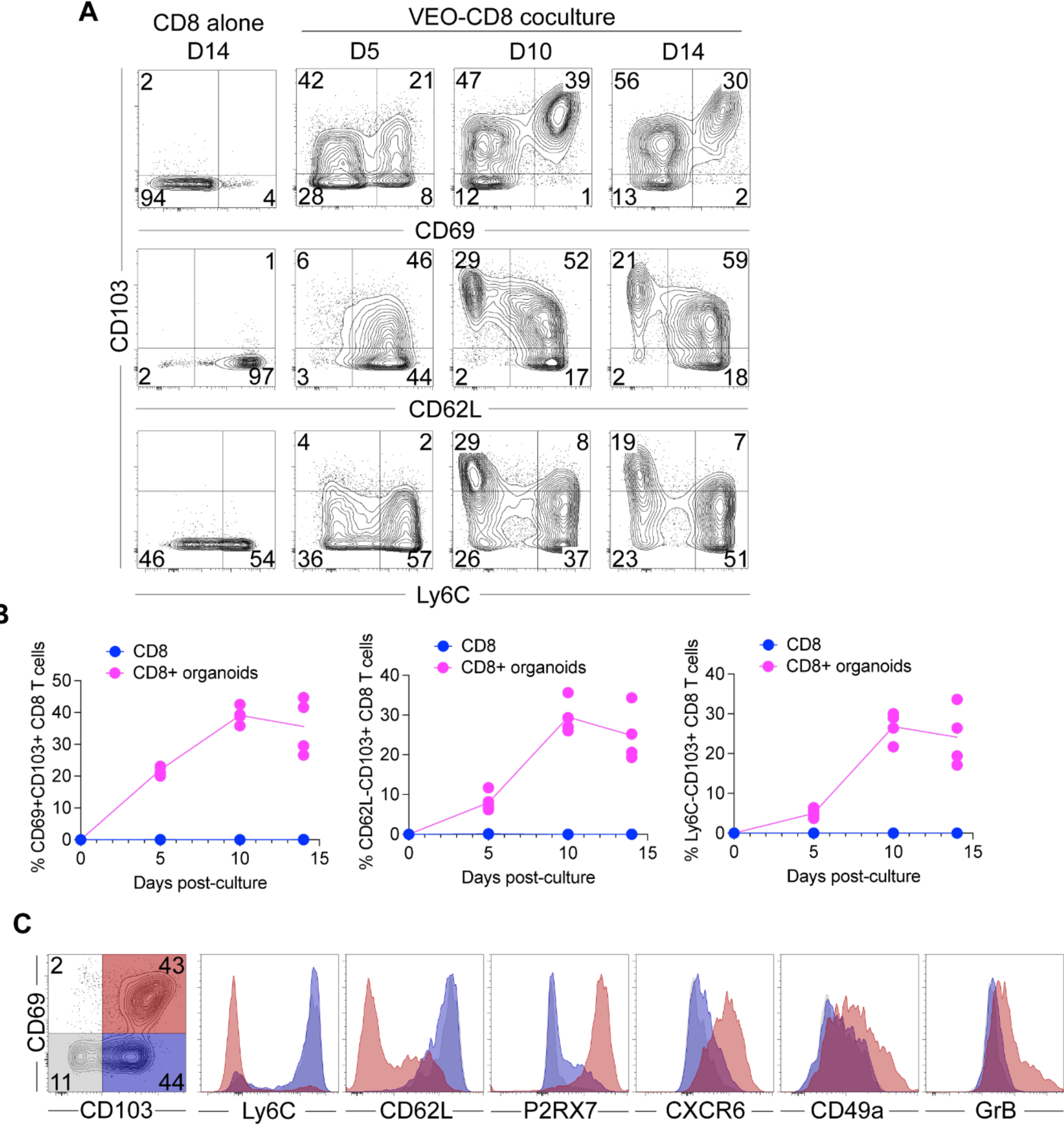
The induction of TRM phenotype by the VEOs is a gradual process, and among various subsets generated post-co-culture, the CD103+CD69+ subset alone phenotypically resembles true epithelial TRM. A. Phenotype of CD8 T cells cultured with VEOs for indicated time points was assessed by flow cytometry. The left most column represents cells maintained in the absence of VEOs for 14 days. Representative flow plots depicting expression of various TRM-associated markers are shown. Cells were gated on live and congenic marker (CD45.1 or CD90.1). B. Scatter plot depicting percentage of CD8 T cells positive for various TRM-associated markers across time. C. Flow-based comparison of TRM markers among the 3 subsets generated by the co-culture (CD69+CD103+, CD69-CD103+ and CD69-CD103-) showing that only the phenotype of CD69+CD103+ population aligns with previously described TRM phenotype including CD62L-, P2rx7+, CD49a+, CXCR6+ and a fraction of cells that are granzyme-B+. Data are representative of two repeats with n=4-6/condition.

### Transcriptional alignment of *in vitro* generated TRM with bonafide *in vivo* TRM

The phenotypic resemblance between VEO-induced CD8 TRM and CD8 TRM established *in vivo* upon viral infection strongly suggests that the *in vitro* generated CD69+ CD103+ CD8 T cells faithfully resemble TRM. However, a number of these phenotypic markers can arise during T cell activation and cytokine stimulation and have led to questioning the establishment of TRM identity by phenotyping alone. Detailed transcriptional analysis of TRM across tissues and species have established a core-TRM transcriptional signature that has been used to establish the fidelity and identity of particular TRM populations ^29,36^. We performed population-based RNA-seq analysis comparing the CD69+CD103+ *ex vivo* generated CD8 T cell subset with CD8 T cells maintained without the VEOs. Out of the 6,223 (3,748 up- and 2,475 downregulated) differentially expressed genes between the 2 cell types, many of the top 25 up- and downregulated genes (e.g. upregulated: *Gzma*, *Hic1, Ccr9, Itgae*; downregulated: *Eomes*, *Sell*, *Klf2*, *S1pr1*) are similarly regulated in bonafide TRM (Fig. 3A) ^11,13,37^. To further substantiate this overlap, we compared the expression of a selected list of genes associated with TRM signature belonging to various T cell-associated processes. A heatmap depicting the expression of these genes between co-cultured CD8 T cells and CD8 T cells cultured alone is shown in Fig.3B. Expression of these same genes between bonafide gut TRM and splenic TCM (extracted from GSE 147080) is shown on the right (Fig. 3B).

**Figure 3:**
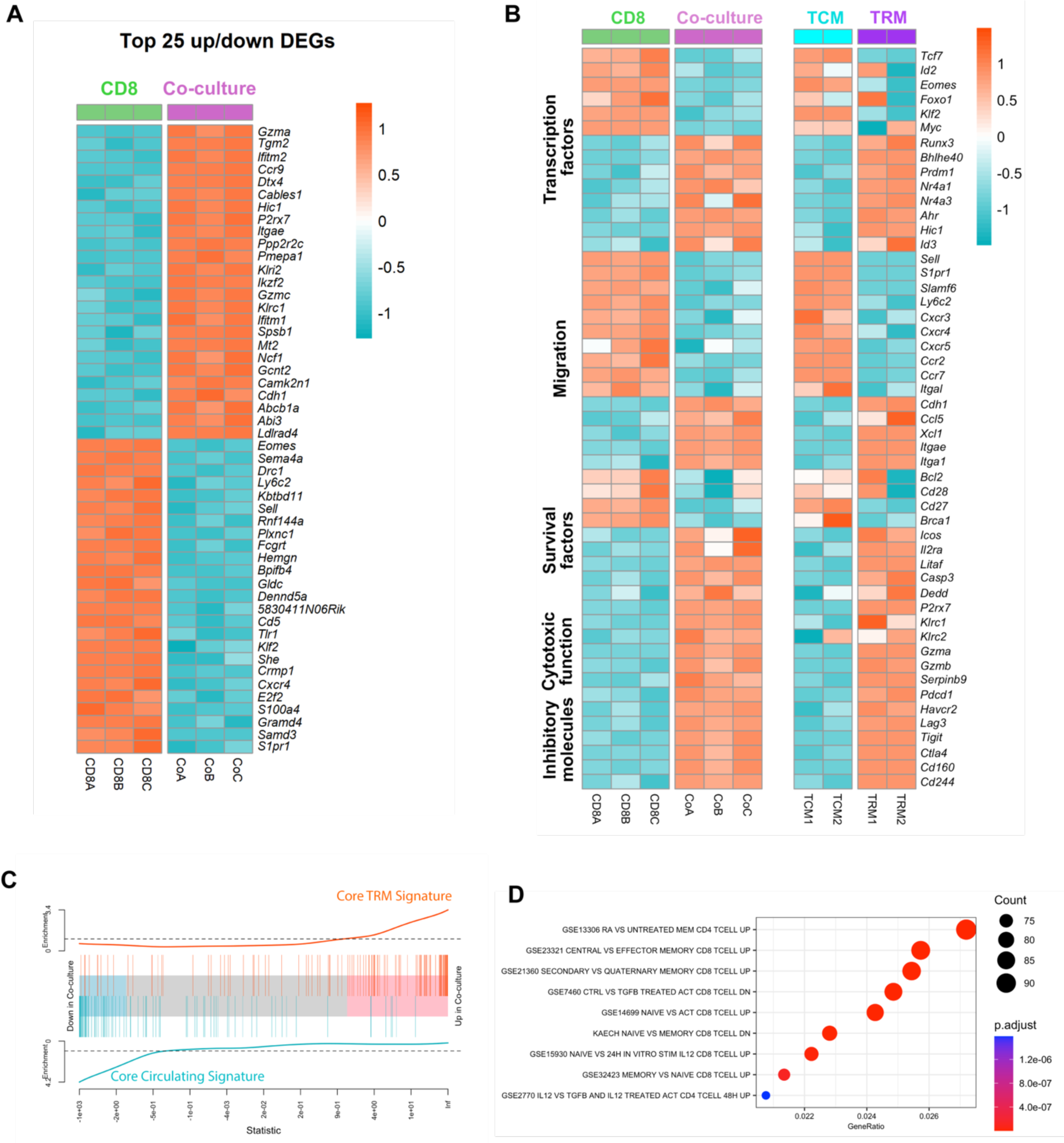
Transcriptional overlap between bonafide in vivo TRM and in vitro differentiated TRM. **A.** Heatmap of top 25 differentially up-/downregulated genes between CD8 TRM generated via co-culture with VEOs vs CD8 T cells maintained alone. The score was calculated as - log10(*padj*)*log2FC, and differentially expressed genes (DEGs) were based on this score. **B.** Expression of selected gene sets belonging to indicated categories between CD8 and co-cultured TRMs (CD69+ CD103+). Expression level of these same genes for circulating CD8 TCM and CD8 TRM from published data set (GSE 147080) is shown on the right. **C.** GSEA plot. Core TRM and core circulating gene signature was created using a ranked gene list from published data comparing TRM and TCM^12^. Enrichment of the overexpressed and underexpressed *in vitro* TRM gene sets in this ranked list is plotted. **D.** Enriched pathways in *in vitro* TRM based on MSigDB are shown.

The *in vitro* TRM exhibited differential expression of a number of transcription factors associated with enforcing residency, such as downregulation of *Tcf7*, *Klf2*, *Eomes* and upregulation of *Runx3*, *Bhlhe40*, and *Prdm1*. Similarly, many of the genes associated with migration (e.g. *Sell*, *Ccr7*, and a number of other chemokine receptors) were downregulated in *ex vivo* TRM as well as *in vivo* established TRM (Fig. 3B). However, integrin and cadherins that help anchor TRM to local tissues (e.g. *Itgae*, *Cdh1* and *Itga1*) were upregulated in both *in vitro* and *in vivo* TRM. Similarly, VEO-induced TRM showed heightened expression of cytotoxic molecules as well as co-inhibitory receptors (the latter help regulate uncontrolled T cell activation). This signature also closely resembled what has already been shown for *in vivo* TRM. A gene set enrichment analysis (GSEA) found significant enrichment of core TRM signature genes (extracted from publicly available data) in the *in vitro* TRM and negative enrichment of genes associated with circulating CD8 T cells (Fig. 3C). A further analysis of biological processes overrepresented in the co-cultured TRM within the MsigDB database showed enrichment of a number of pathways upregulated in the memory CD8 T cells compared to naive (Fig. 3D). Interestingly, we also noticed genes upregulated in response to retinoic acid (vitamin-A, Retinoic Acid/RA) and TGF-β are represented among these pathways. In summary, our transcriptomic analysis showed strong overlap of previously established TRM gene signatures among the *in vitro* generated CD8 TRM, further verifying their TRM identity.

### Reactivated circulating memory CD8 T cells can differentiate into TRM under influence of VEOs

After establishing that VEOs can support differentiation of effector CD8 T cells (generated from activation of naive CD8 T cells) into mature TRM *in vitro* with remarkable efficiency, we tested whether they could also facilitate TRM differentiation of circulating memory CD8 T cells. For this, we first generated central and effector memory CD8 T cells (TCM and TEM) *in vivo* by transferring naive congenically marked P14 CD8 T cells (CD45.1+) to C57Bl/6j mice followed by LCMV infection. The P14 CD8 T cells were allowed to differentiate into circulating memory CD8 T cells for 75 days post-infection, at which point the SLOs from these mice were isolated and TCM (CD45.1+, CD44 hi, CD62L hi) as well as TEM (CD45.1+, CD44 hi, CD62L lo) were separated by flow sorting. Sorted TCM and TEM were co-cultured with VEOs to induce differentiation for 10 days. As a positive control, we also included *in vitro* generated effector CD8 T cells, which have been shown to differentiate into CD69+CD103+ TRM. Although a fraction of TCM and TEM survived in co-culture, they failed to adopt the epithelial TRM phenotype (Fig.4A&B). In contrast, circulating memory CD8 T cells exposed to VEOs loaded with cognate antigenic peptide (gp33) formed TRM-like cells (Fig.4A&B). This suggests circulating memory CD8 T cells require antigenic restimulation to enable their differentiation into TRM. However, the efficiency of adoption of various TRM-associated markers (CD69+CD103+, CXCR6+ and CD62L-) was significantly lower among the reactivated TCM and TEM compared to effector CD8 T cells (Fig.4B). The CD8 TCM showed better acquisition of TRM phenotype compared to TEM, although this difference was not statistically significant. Altogether, this suggests circulating memory CD8 T cells can be programmed into TRM but need reactivation for differentiation.

**Figure-4:**
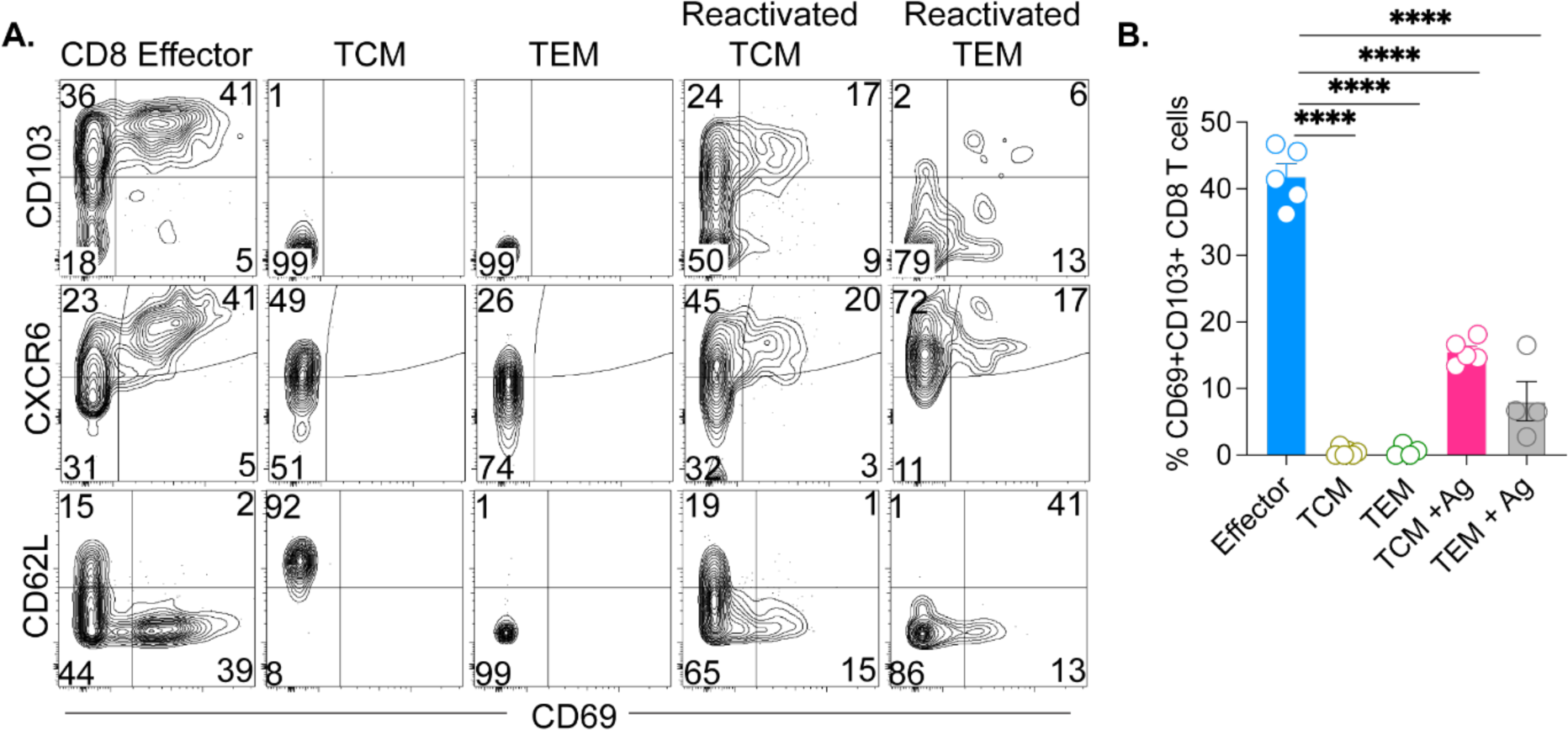
Circulating memory CD8 T cells need to be reactivated to form TRM under the influence of VEOs. C57Bl/6j mice received 104 CD45.1+ naïve P14 CD8 T cells and were infected with LCMV. At 70 dpi, SLOs were harvested, and TCM (Live CD8a+ CD45.1+ CD62L+) and TEM (Live CD8a+ CD45.1+ CD62L-) cells were flow sorted and incubated with VEOs for 10 days. In some cases, the cells were exposed to epithelial cells loaded with gp33 peptide (0.2 μg/ml) labeled as reactivated cells. Naïve CD8 T cells differentiated *in vitro* and co-cultured with VEOs were included as a control (effector). Representative flow plots are shown in **A**, gated on live congenic marker (CD45.1) T cells, and percentages are enumerated in **B**. Data is representative of one repeat with n=5/condition. Bars indicate mean ± SEM. One-way ANOVA with Holm-Sidak multiple comparison test (B). ****<0.0001.

### VEO-induced CD8 TRM remain functional and can be generated in the absence of physical contact with the organoids

TRM located in frontline mucosal tissues rapidly elicit cytotoxic granules and cytokines after TCR stimulation, and maintenance of this functionality is crucial to limit pathogen replication. Here we assessed whether the *in vitro* generated CD8 TRM remain functional in response to antigenic recall. For this, wells containing CD8 TRM and VEOs (14 days post-co-culture) were treated with antigenic peptide in the presence of Brefeldin-A, and the expression of various cytokine molecules were checked by intracellular cytokine staining followed by flow cytometry. As shown in Fig. 5A&B, in response to the peptide challenge CD8 T cells elaborated significant amounts of IFN-γ, TNF-α and IL-2. These data indicate TRM generated in response to VEO-derived cues retain their functional potential as has been shown for *in vivo* TRM^9^.

**Figure-5:**
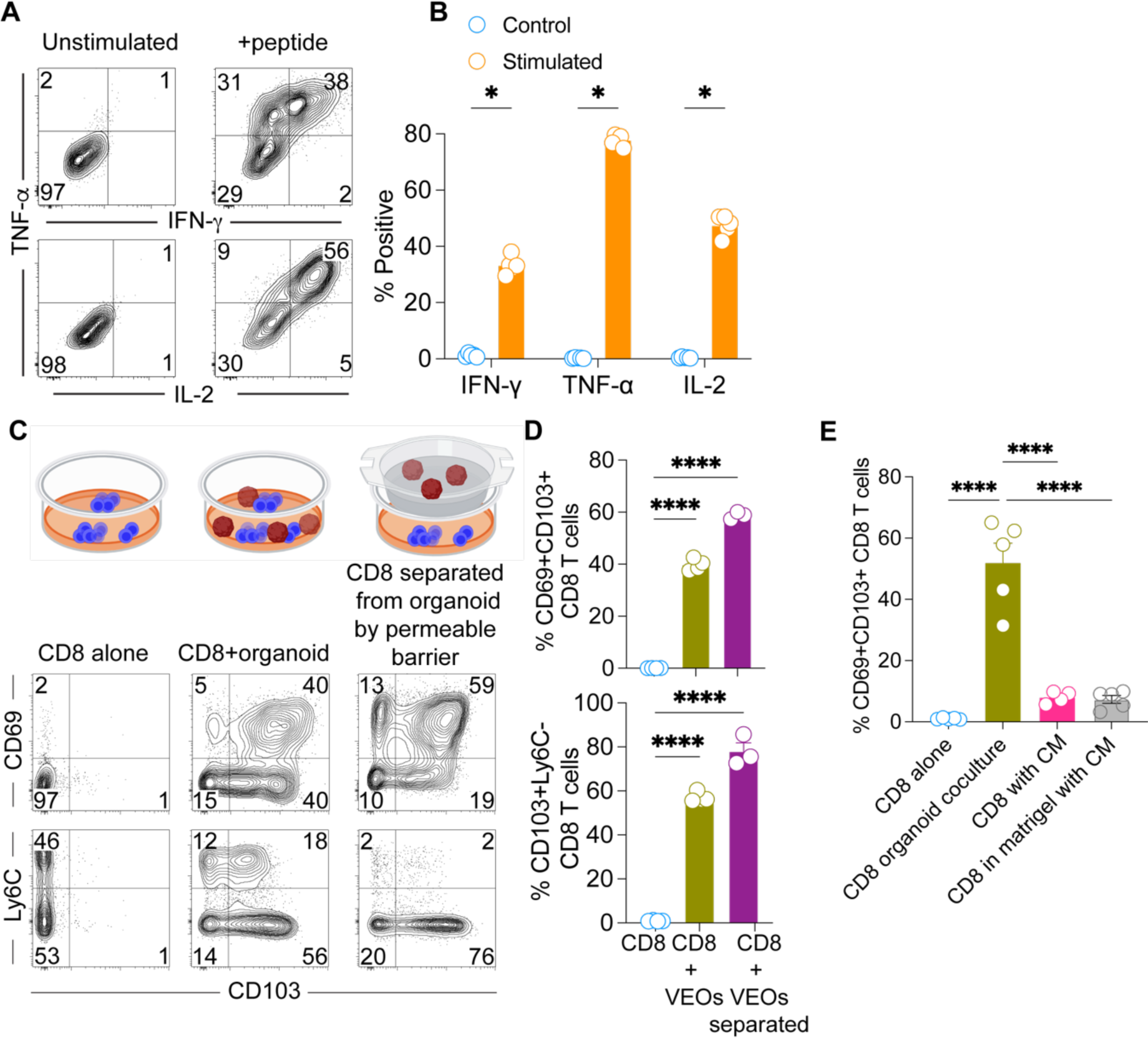
In vitro differentiated CD8 TRMs remain functional and could be generated in the absence of physical association with VEOs. **A**. Co-cultured CD8 T cells (Day 11) were stimulated with antigenic peptide or unstimulated for 4 hours in the presence of Brefeldin-A. Representative flow plots showing expression of cytokines IFN-ψ, TNF-α and IL-2 are shown. Plots are gated on live CD8β+ T cells. **B**. Percentage of stimulated cells expressing various cytokines are compared against unstimulated cells. **C**. Transwell assays were conducted whereby CD8 T cells in the bottom chamber were exposed to soluble mediators released from VEOs for a period of 10 days. This was compared to CD8 T cells cultured in the absence of VEOs and CD8 T cells embedded together with VEOs. Representative flow plots are gated on live CD8β+ T cells showing robust adoption of TRM phenotype when CD8 T cells were separated from VEOs by the semipermeable barrier. **D**. Bar graph showing percent positivity of various TRM phenotypes. **E**. Regular exposure to VEO-conditioned media (CM) for 10 days was not sufficient to drive CD69+ CD103+ epithelial TRM phenotype. Bar graph comparing various CM treatments with the regular co-culture system is shown. Data are representative of three repeats with n=3-6/condition (B, D), and two repeats with n=5/condition (E). Bars indicate mean ± SEM. Multiple Student’s t-tests (B) One-way ANOVA with Holm-Sidak multiple comparison test (D, E). * < 0.05, ****<0.0001.

Next, we tested if the *in vitro* TRM differentiation process relies on direct interaction with VEOs or can be achieved when the VEOs and CD8 T cells are physically separated. For this we used transwell inserts containing permeable membranes such that the CD8 T cells can access any soluble factors produced by the VEOs but are not in direct contact. We exposed the CD8 T cells to VEOs through the permeable barrier for up to 15 days and evaluated the CD8 T cell phenotype by flow cytometry. Interestingly, these CD8 T cells upregulated the classical TRM markers CD69 and CD103 (Fig. 5C&D). For comparison, we also had wells without the transwell inserts where CD8 T cells were either maintained alone or embedded in the VEOs co-culture system. As expected, the co-cultured CD8 T cells upregulated TRM-associated markers. These results showed that TRM differentiation can be mediated by the soluble factors produced by epithelial organoids. We next asked whether induction of TRM phenotype can be achieved through regular supplementation of conditioned media (CM) from wells containing VEOs. Exposure of effector CD8 T cells to VEO-derived CM (every 2 days for 10 days) did not dramatically upregulate CD69 or CD103 expression (Fig.5E). Embedding the CD8 T cells in BME also failed to induce a TRM phenotype. Altogether, these data suggest that while *in vitro* TRM differentiation can be induced by soluble agents, these factors might be labile in nature and require continuous contact with responding CD8 T cells.

### VEOs support viral replication, and the organoid co-culture system can be used to probe molecular drivers of antiviral TRM differentiation

The vaginal epithelium is a common portal for viral invasion and often serves as an initial replication site before the pathogen spreads to distal organs. As such, understanding the viral replication dynamics in the vaginal mucosa and the ensuing immune response is crucial to improve antiviral therapies and vaccines. Here, we aimed to test whether VEOs can be targeted by a common sexually transmitted infection, Herpes Simplex virus (HSV). For this, fully grown VEOs (>7 days post subculture) were released from the BME through depolymerization of the extracellular matrix and were exposed to a recombinant HSV-1 K26GFP that encodes a green fluorescent VP26 capsid protein^38^. Infected cells showed punctate green signals, which correspond to capsid assembly sites within the nucleus between 24-36 hours post-infection (Fig.6A)^38^. These findings suggest VEOs can support HSV replication and could be used to test protective efficacy of drugs and immune cells in a more physiological setting than what is afforded by routine *in vitro* cell culture models.

**Figure-6:**
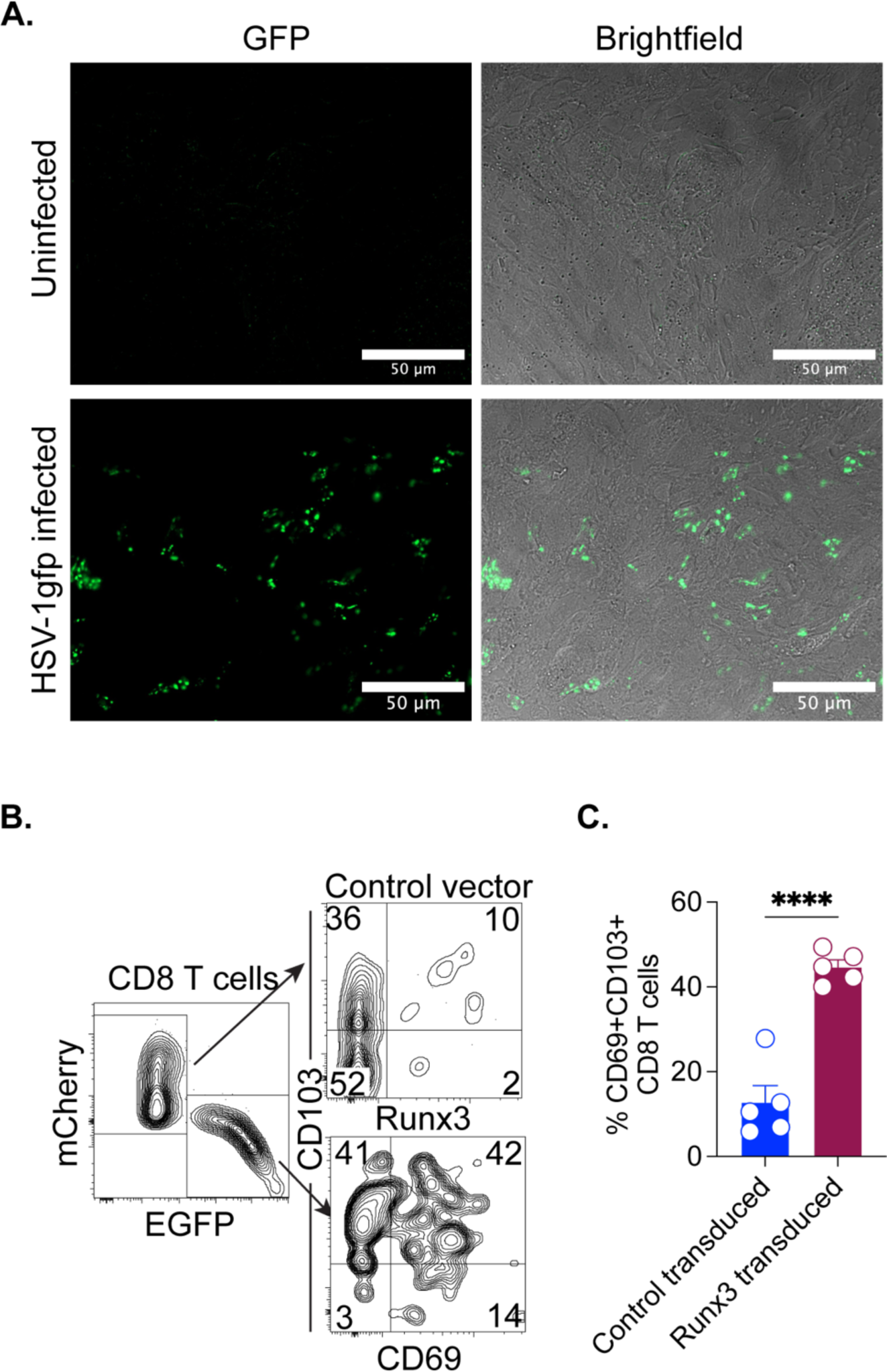
VEOs support viral replication, and the co-culture system is amenable to genetic perturbation. **A**. Seven-day old VEOs were released from basement membrane extract and were maintained as a suspension culture for 24 hours. Then they were infected with HSV-1K26GFP or mock infected. Wells containing infected and uninfected cells were visualized 36-hours post-infection using fluorescence microscopy. Representative images are shown. Scale bar-50 μm. **B**. *In vitro* activated P14 CD8 T cells were retrovirally transduced with Runx3-EGFP expressing vector or control-mCherry expressing vector. Equivalent number of cells were cultured with VEOs for 10 days, and their ability to form TRM was tested by flow cytometry. Representative flow plots of total transgene positive P14 cells are shown in the left, and the level of CD69 and CD103 on the two reporter positive populations are shown in the right. **C**. Bar graph comparing percentage of CD69+ CD103+ cells among the two transduced populations. Bars indicate mean ± SEM. Student’s t-test. ****<0.0001

Despite the outsized functional role of TRM cells in infection, the molecular drivers coordinating TRM fate remain ill-defined. Most studies utilize rodent models to test the role of putative regulators, an approach that is not often scalable and technically challenging. We wanted to test if the *in vitro* VEO-mediated CD8 TRM differentiation system could be used to define regulators of TRM fate. Notably, we detected elevated expression of the transcription factor Runx3 in VEO-co-cultured CD8 T cells compared with the CD8 T cells alone (Fig.3B). Runx3 has also been established as a key transcription factor that promotes TRM formation in the intestine^12^. However, the role of Runx3 in governing TRM fate in the FRT remains unknown. To test whether Runx3 influences FRT TRM formation, we transduced activated CD8 T cells with either a Runx3 encoding retrovirus (simultaneously encoding an EGFP reporter) or a control vector encoding mCherry. Transduced CD8 T cells were mixed at 1:1 ratio and were co-cultured with the VEOs for 5-10 days. We found a significantly higher percentage of CD69+CD103+ TRM cells among the Runx3 transduced cells compared to the control vector (Fig.6B&C). These data suggest Runx3 drives FRT CD8 TRM formation, and more importantly, our findings establish a proof of principle that the VEO-CD8 co-culture system can be used to identify molecular regulators of TRM differentiation.

A critical advantage of the *in vitro* differentiation system is the generation of an abundant (near unlimited) number of TRM compared to the sparse number of TRM that can be isolated from the FRT *in vivo* ^39^. Comparison of the relative TRM yield between the two systems showed that a single well of a 96 well plate could generate ∼3 times more CD8 TRM than what could be extracted from a single mouse lower FRT that was infected with LCMV intravaginally 30 days before (Supplementary Fig.3). Altogether, these attributes establish the robustness of the VEO-CD8 co-culture model for studies of antiviral TRM differentiation and function.

**Supplementary Figure 3.**
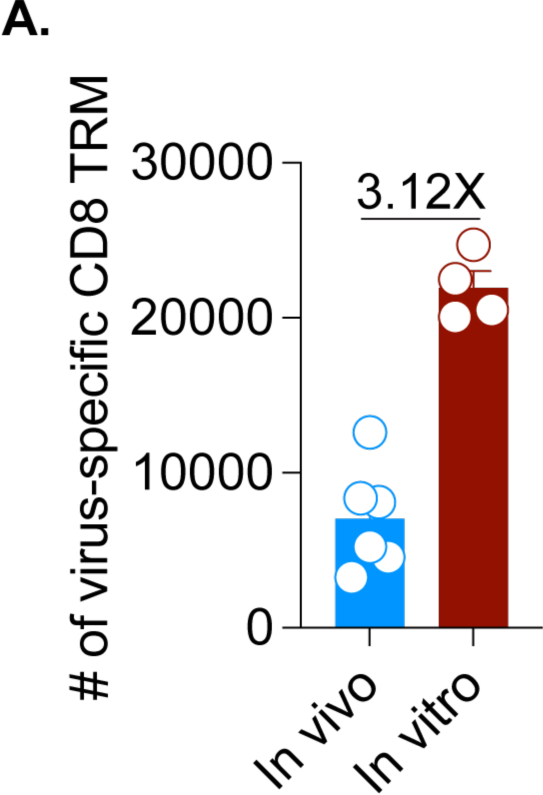
Comparison of FRT TRM cell yield between *in vivo* mouse model of intravaginal LCMV infection (infected 50 days prior) and VEO-induced *in vitro* TRM isolated from a single well of a 96-well plate. Data are representative of two repeats with n=4-6/condition. Bars indicate mean ± SEM. Student’s t-test ****<0.0001

### Inhibition of TGF-β signaling impairs TRM differentiation in organoids

TGF-β is a multifunctional cytokine has been implicated in epithelial TRM differentiation in numerous tissues ^24,26,40,41^. Our pathway analysis also showed an important role of TGF-β in programming TRM differentiation in the VEO-CD8 co-culture system (Fig.3D). Consequently, we aimed to test the relevance of TGF-β signaling in *in vitro* vaginal TRM differentiation using two separate approaches. In the first approach, we used two distinct pharmacological inhibitors that block separate aspects of TGF-β signaling. The small molecule SB431542 is a potent and selective inhibitor of TGF-β type-1 receptor kinase (ALK-5) but also affects ALK-4 and ALK-7 ^42^. Treatment with SB41532 is thought to inhibit signaling through the TGF-β receptor. When co-cultured CD8 T cells were treated with 10 μM of SB431542 for 7 days, it led to an almost complete absence of CD69+ CD103+ TRM cells (Fig. 7A&B). This treatment also led to the CD8 T cells failing to downregulate CD62L, a cardinal feature of TRM (Fig. 7A&C). Next, we tested another small molecule inhibitor, CWHM-12, which specifically targets ɑV integrins ^43^. AlphaV integrin mediated processing of inactive TGF-β to active TGF-β has been shown to be important for CD8 TRM formation ^44–47^. VEO-CD8 co-cultures were treated with various concentrations of CWHM-12, which led to a dose-dependent reduction in the percentage of CD69+CD103+ TRM (Fig. 7D&E). However, another property of epithelial TRM, i.e. downregulation of Ly6C expression, was not altered in CWHM-12 treated cells (Fig. 7D&E). Altogether, our results suggested that *in vitro* FRT TRM differentiation could be prevented by pharmacological inhibition of TGF-β pathways.

**Figure-7:**
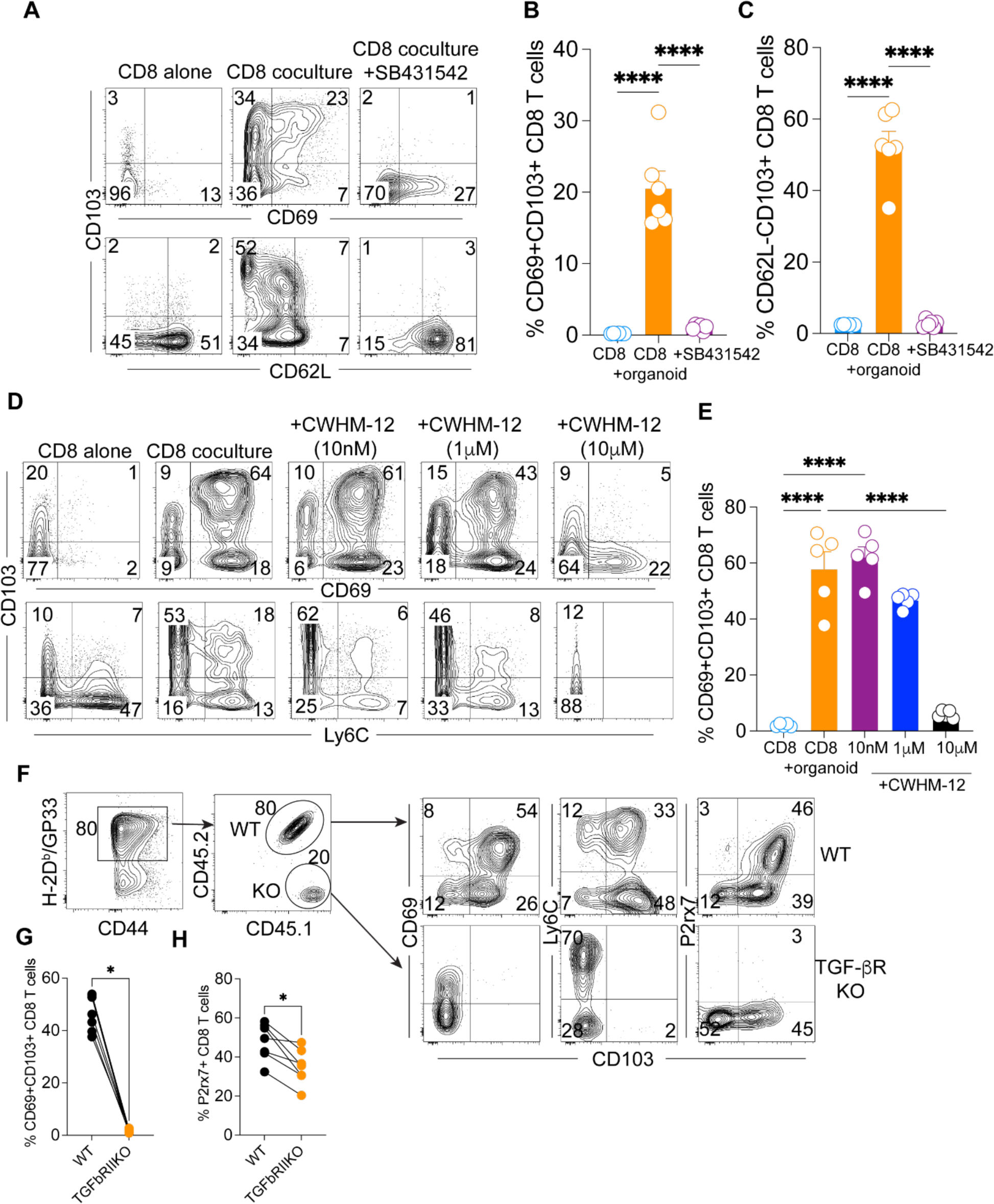
Pharmacological and genetic inhibition of TGF-β signaling interferes with in vitro TRM generation. **A**. CD8 T cells co-cultured with VEOs were treated with TGF-β signaling inhibitor, SB431542 (10 µM), or vehicle control for 7 days. Representative flow plots of live CD8β+ T cells are shown. **B, C**. Percentage of TRM phenotype cells are enumerated. CD8 T cells maintained alone are included as a control. **D**. CD8 T cells co-cultured with VEOs were treated with increasing concentrations of an inhibitor of TGF-β activating ɑv, CWHM-12 (10 nM to 10 µM), or vehicle for 12 days. Representative flow plots of live CD8β+ T cells are shown. **E**. Percentage of CD69+CD103+ phenotype cells are enumerated. **F**. TGF-β receptor deficient CD8 T cells fail to adopt TRM phenotype in the VEO co-culture model. Wild type (WT, CD45.1+CD45.2+) and TGF-βRII deficient P14 (KO, CD45.1+CD45.2-) CD8 T cells were activated and embedded in BME containing VEOs at 1:1 ratio. Representative flow plots 12 days after the co-incubation are shown. Total P14 CD8 T cells and the ratio of WT and KO CD8 T cell percentages retained as well as their associated phenotypes after 12 days are shown. **G**. Comparison of CD69+CD103+ CD8 T cells and **H**. p2rx7+ CD8 T cells between WT and KO groups. Data are representative of two repeats with n=4-6/condition. Bars indicate mean ± SEM. One-way ANOVA with Holm-Sidak multiple comparison test (B, C, E). Wilcoxon matched-pairs signed rank test (G, H). * < 0.05, ****<0.0001

To further substantiate the role of TGF-β signaling in the FRT TRM differentiation process, we used a genetic approach. We used a previously described genetic model system where dLck-cre mice were crossed to *Tgfbr2* flox mice, permitting conditional depletion of TGFβRII expression in mature T cells. Transgenic P14 CD8 T cells from TGF-βR conditional knockout (KO) donors and their wild type (WT) counterparts were enriched and activated *in vitro* before being introduced to the VEO-CD8 co-culture system at 1:1 ratio. Twelve days after the co-culture, we performed phenotypic analysis of the resulting T cell population by flow cytometry. As shown in Figure 7F, among all the antigen-specific CD8 T cells (H2-Db:gp33 tetramer positive) present, the KO CD8 T cells were present at approximately 4-fold lower rate than their WT counterpart. The KO CD8 T cells also failed to upregulate CD103 (Fig. 7F&G). Lack of TGF-β receptor signaling also impaired CD8 T cells’ ability to downregulate Ly6C and upregulate P2rx7, the latter of which is a known TGF-β regulated gene in TRM ^48^. Altogether, this experiment further supports the crucial role of TGF-β signaling in mediating TRM differentiation in the VEO system.

## Discussion

Mice have long been the model of choice in fundamental TRM studies and have contributed immensely to our understanding of TRM biology. However, a number of issues with the *in vivo* model have restrained progress in generating a comprehensive picture of TRM differentiation. Chief among these is the highly inefficient extraction of TRM from tissues via enzymatic digestion; by performing microscopy-based counting, Steinert et al. found enzymatic extraction could only isolate 1 CD8 TRM cell for every 69 actual CD8 TRM cells present in the FRT ^39^. This ratio is less biased for other tissues like the small intestine (12.9) and liver (6.13) but is nonetheless widespread. Moreover, there was bias in the extraction of cells bearing different phenotypes e.g., CD103+ TRM were extracted more easily than the CD103-TRM ^39^. Recent work has also suggested that the routine enzymatic digestion processes can alter the transcriptome of isolated cells potentially leading to confounding results ^49^. TRM are also highly susceptible to cell death upon isolation, complicating phenotypes and outcomes ^50–52^. Lastly, separating the tissue-specific signals responsible for local TRM differentiation from systemic signals that impact other linked processes like initial T cell activation, migration and entry into the NLT is difficult in mouse models. Here, we sought to address these limitations by establishing a robust *in vitro* system for modeling TRM differentiation with epithelial organoids that solely focuses on local TRM differentiation under the influence of inductive cues produced by NLTs. We demonstrated that the vaginal epithelium alone is sufficient to orchestrate CD8 TRM differentiation. Importantly, our approach establishes a path for the development of reductionist, *in vitro* immune-epithelial co-culture models to interrogate aspects of biology that can’t be efficiently modeled *in vivo*. This opens avenues for conducting TRM-centric functional, genomic, and metabolomic assays that requires higher cell numbers than routine flow cytometry and transcriptional methods.

Despite its reductionist nature, the VEO system faithfully recapitulates the stratified squamous epithelium of the *in vivo* vaginal tissue, which is made up of basal, suprabasal, and cornified apical epithelium. Moreover, single cell RNA-sequencing analysis has revealed at least 6 separate clusters of transcriptionally distinct epithelial cells among these layers of vaginal epithelium ^20^Our co-culture system exposes CD8 T cells to products of each of these distinct epithelial cell types which is hard to model in classical immortalized vaginal epithelial cell lines that cannot differentiate into these various cell types. Moreover, the VEO co-culture model enables the detailed characterization of events specifically occurring at the final site of TRM residence, circumventing confounding factors present in live animal studies, such as the impact of CD8 T cell entry into NLT stroma on tissue residence. It is noteworthy that our model, while not incorporating the vaginal microbiome, offers a platform amenable to introducing bacterial species or their metabolites, enabling a detailed examination of a tripartite interaction involving epithelium-commensal microbiome-immune cells. There is a significant gap in our understanding of the impact of the vaginal microbiome on adaptive immunity, and our system could be used to fill this need.

Previous work has implicated cytokines TGF-β, IL-33, and TNF-α as crucial modulators of CD8 TRM differentiation ^6,13,24^, and our study also suggested that activated CD8 T cells can differentiate into TRM by soluble factors in the absence of physical interaction with epithelial cells. However, an interesting finding from our co-culture studies is the pivotal role of a second antigenic exposure in further enhancing the TRM phenotype. This is in agreement with past studies that have shown the enhancement of CD8 T cell effector response as well as improvement in TRM density with second antigenic exposure^34,35,53,54^. Altogether, it suggests that while cell-cell interaction might not be essential, it greatly improves epithelial TRM density. By utilizing pharmacological and genetic means, we verified the significant contributions of TGF-β. Future investigations will employ high-throughput proteomic screening to unravel the involvement of other proteins in this intricate process.

We showed that VEOs support HSV-1 replication, and as such, this model could be easily adopted for high-throughput screening of drugs or cell-based therapies that will target viral infections of the lower FRT. CD8 T cells in the co-culture model could also be genetically modified using shRNA or CRISPR to delineate the molecular underpinning of TRM development. We provided a proof of principle experiment showing the relevance of Runx3 in FRT TRM development, but this could easily extend to libraries of transcription factors, epigenetic modifiers, and other molecules implicated in T cell biology. Beyond the scientific advancements, our organoid model aligns with ethical considerations in animal research, adhering to the principles of Replacement and Reduction outlined by Russell and Burch in1958 ^55^. By offering an ethically sound alternative to live animal studies, our model not only replaces the need for animal studies with a cell culture approach but also reduces the number of animals required for experimentation.

In summary, our *in vitro* TRM generation system provides a reductionist, scalable alternative that will allow deeper interrogation of TRM biology than what is possible with existing *in vivo* approaches without sacrificing the complexity of the epithelial environment. Our findings suggest that the type-II mucosa-derived signals are sufficient for TRM differentiation and TGF-β is important in this differentiation process. On a broader scale, this approach presents a valuable tool for future exploration into mechanisms that govern immune defense against sexually transmitted infections and other pathogens affecting the FRT.

## Method

### Mice and Infection

C57BL/6j (B6) (strain-000664), CD45.1 mice (strain-033076), CD90.1 mice (strain-000406), and OT-I CD8 T cell transgenic mice (strain-003831) were procured from the Jackson Laboratory and housed at Brown University, Providence, RI. P14 and gBT-I CD8 T cell transgenic mice were kind gifts from David Masopust (University of Minnesota) and Gregoire Lauvau (Albert Einstein College of Medicine) respectively. Congenically marked P14, gBT-I and OT-I genotype mice were generated through crossbreeding original transgenic lines with congenic marker bearing mice strains. The *Tgfbr2*^f/f^ dLck-cre+ and *Tgfbr2*^f/f^ dLck-cre-P14 mice have been described before and were maintained at the animal facility at University of Texas Health at San Antonio (San Antonio, TX). Mice aged between 6-20 weeks were utilized in all experiments, adhering to the guidelines set forth by Brown University’s or University of Texas Health Science Center at San Antonio’s Institutional Animal Care and Use Committee guidelines. Lymphocytic choriomeningitis virus (LCMV)-Armstrong was intravaginally or intraperitoneally administered using 10 uL or 200 uL of sterile RPMI-1460 media containing 2×10^5^ plaque-forming units (PFU), respectively. For intravaginal infections, Depo-provera (3 mg/mouse diluted with sterile PBS) was given subcutaneously 5 days before viral delivery to improve infection efficiency.

### Chemicals, cytokines and peptides

Most chemicals for organoid cultures (EGF, Y-27632 (ROCK inhibitor), and SB-431542) were obtained from Sigma Aldrich. The αV integrin inhibitor, CWHM-12, was synthesized at Washington University, St. Louis and obtained via a collaboration with Peter Ruminski. Recombinant interleukin-2, 12 were purchased from Biolegend. Peptides were synthesized by Alan Scientific to at least 95% purity.

### Establishment of VEO-CD8 co-culture model

For establishing epithelial organoids from murine vaginal tissues, we followed a recently described protocol by Ali et al. ^20^. Briefly, B6 mice aged at least 8 weeks were euthanized, and vaginal epithelium was separated from underlying stroma after overnight digestion with pronase and DNaseI. A single cell suspension of vaginal epithelial cells was prepared by pipetting, mixed with Cultrex™ Basement Membrane Extract (BME) (RnD Systems), and plated in 24 well plates with organoid culture medium (OC) containing DMEM/F12 media supplemented with the following agents: 1% Penicillin/streptomycin, 0.2 ug/mL Amphotericin B, 2% B27 Supplement, 5 μM SB431542, 100 ng/mL murine Epidermal growth factor (EGF), and 10 μM Y-27632 added for the first 4 days of culture. Epithelial stem cells were allowed to differentiate and form circular organoids for 7-14 days, at which point further subculturing was done to propagate the VEOs. For most co-culture studies, VEOs between passage number-3 and -8 were used. Spleen and lymph nodes from C57Bl/6j mice were isolated after euthanasia, and naïve CD8 T cells were isolated using a magnet-based negative enrichment protocol following the manufacturer’s direction (Mojosort mouse CD8 naïve T cell isolation kit, Biolegend). These CD8 T cells were activated in the presence of anti-CD3ε (Biolegend), B7-1Fc (Biolgened), IL-2 (10 U/ml) and IL-12 (2.5ng/ml) for 2 days. Afterwards, the expanded CD8 T cells were transferred to a new 24 well plate and rested for 2 days with IL-2 alone (10U/ml). Then the effector CD8 T cells were mixed with epithelial cells obtained from trypsinized VEOs, and the cell mixture was resuspended in BME and plated at 8 μL per well on a 96-well plate. Following a 30-minute incubation upside down at 37°C, 200 μL of T cell-OC culture medium (T/OC) (1% Penicillin/streptomycin, 0.2 ug/mL amphotericin B, 2% B27 Supplement, 2mM L-glutamine, 1% non-essential amino acids, 1% sodium pyruvate, 55 uM Beta-mercaptoethanol, 100 ng/mL EGF, 10 U/mL IL-2, and 10 uM Y-27632 added for the first 4 days of culture) media was added to each well. Media changes occurred every two days during the culture, ensuring careful handling to preserve T cells lodged in the plate.

### Transwell experiment

Transwell experiments utilized MatTek cell culture inserts with 0.4 μm membranes. Organoids were trypsinized, quenched with 10% FBS in RPMI, and washed in T/OC media. The resulting single-cell suspension was either directly plated on the insert in 50 μl BME or combined with activated CD8 T cells before being plated in the bottom well at a density of 200,000 CD8 T cells per 50 μL Matrigel. After a 30-minute upside-down incubation at 37°C, 500 uL of T/OC media were added to each well. In the wells where VEOs and CD8 T cells were present in separate chambers, approximately 300,000 effector CD8 T cells were added to the lower chamber. Media in the bottom well was changed every two days. After incubation, T cells in the lower chamber were analyzed, unless otherwise stated.

### Lymphocyte isolation and phenotyping

For lymphocyte isolation from *in vitro* cultured cells, well contents were collected and washed in PBS, and cells were used for staining. The lymphocyte isolation from secondary lymphoid organs (SLOs) and non-lymphoid tissues (NLTs) was performed as described with small modifications^56^. Lymphoid tissues were mashed using the plunger of a 3-mL syringe and filtered through 70 μm mesh before staining. Female reproductive tract tissues were chopped into small pieces and incubated with RPMI+2.5% FBS containing collagenase type-IV (Sigma, 1mg/ml) and Dnase I (Sigma, 2μg/ml) at 37°C with constant shaking for 45 min. After the incubation, tissues were further dissociated using a gentlemacs dissociator (Miltenyi Biotec) and filtered twice through a 70 μm mesh before staining.

Isolated lymphocytes were surface-stained with antibodies against CD8α (53-6.7), CD8β (YTS156.7.7), CD45.1 (A20), CD90.1 (OX-7), CD45.2 (104), CD62L (MEL-14), CD44 (IM7), CD69 (H1.2F3), CD103 (M290 or 2E7), Ly6C (HK1.4), CD49a (Ha31/8), PD1 (RMP1-30), P2rx7 (1F11), Epcam (G8.8), and CXCR6 (SA051D1). The following intracellular targets were also detected using antibodies-IFN-γ (XMG1.2), TNF-α (MP6-XT22), IL-2 (JES6-5H4), Tbet (4B10), Eomes (Dan11mag), granzyme-B (QA16A02), and Ki67 (B56). The above antibodies were purchased from Biolegend, BD Biosciences, or Invitrogen. Cell viability was determined using Ghost Dye 780 (Tonbo Biosciences). For intracellular transcription factors and granzyme-B, the Tonbo Transcription factor staining buffer set was utilized. For intracellular cytokine staining after restimulation, the BD Cytofix/Cytoperm kit was used. Antigen-specific CD8 T cells were detected by staining with tetramers (gp33 tetramer for P14, SL8 tetramer for gBT-I, or SIINFEKL tetramer for OT-I) conjugated to brilliant violet-421 dye obtained from the NIH tetramer core facility. The stained samples were acquired using Aurora spectral cytometer (Cytek) and analyzed with FlowJo software (Treestar).

### Confocal Immunofluorescence Microscopy

VEOs or co-culture systems were plated in a chambered cell culture slide with 50 μL of BME per well. T/OC or OC media (500 μL) was changed every two days. After 6-14 days, each sample was fixed (60 minutes at room temperature in 4% Paraformaldehyde) and blocked/permeabilized (overnight at 4°C in 5% normal donkey serum/0.5% Triton X-100/1X PBS). Samples were stained with unconjugated polyclonal rabbit anti-Ki67 (Abcam), unconjugated polyclonal rabbit anti-keratin-5 (Biolegend), Phycoerythrin conjugated anti-Epcam monoclonal (G8.8, Biolegend), and Phycoerythrin conjugated anti-CD90.1 monoclonal (OX-7, Biolegend). Donkey anti-rabbit Cy3 conjugated antibody (Jackson Immunoresearch) was used as a secondary antibody. Primary antibodies were incubated at 4°C overnight, whereas secondary antibody was used at room temperature for 1-1.5h. DAPI was used to visualize the nucleus. Samples were washed with PBS between each step. Slides were mounted with ProLong Diamond Antifade (Invitrogen) before being imaged on an Olympus FV3000 Confocal Microscope. Captured images were processed in Fiji for visualization.

### RNA-seq and analysis

RNA was extracted from CD8 T cells using the RNeasy Plus Micro Kit (Qiagen), and libraries were constructed and sequenced on Illumina NovaSeq 2 × 150 bp paired-end sequencing (Novogene). Adapter sequences and low-quality sequences were trimmed from the raw sequence reads using Trimmomatic v0.36 ^57^. STAR v2.7.3a ^58^ was used to align the trimmed sequences to the mm10 mouse genome and to estimate the number of reads per gene. Gene count was normalized and differentially expressed genes (DEGs) were identified if Padj < 0.05 in DESeq2 v1.38.1 ^59^. Enrichment pathway analysis utilized upregulated genes in co-cultured samples and was performed with ClusterProfiler v4.7.1 ^60^ using MSigDB ^61^. Previously published gene lists for core TRM and circulating signature^12^ were used for Gene Set Enrichment Analysis (GSEA), and it was performed and visualized using the limma v3.54.1 The gene expression pattern of CD8 and co-cultured samples was compared to the previously published TCM and TRM signatures dataset GSE 147080 ^62^ and visualized using pheatmap v1.0.12.

### Quantitative PCR (qPCR)

VEOs were harvested at post-culture days 5, 9, and 15, followed by resuspension in 1 mL of cold 5 mM EDTA in DPBS in 1.5 mL Eppendorf tubes. Subsequently, the suspension was incubated on ice for 30 min. After incubation, samples were washed in 1 mL of cold DPBS by centrifugation at 1,000 x g for 5 min at 4 °C, repeated twice. For the final wash, samples were collected at 1,200 x g for 5 min at 4 °C. The resulting pellets were resuspended in 1 mL of TRI Reagent (Zymo Research) and incubated for 5 min at RT. 0.2 mL of chloroform was added to the tube, and the tube was shaken vigorously followed by 5 min incubation at RT. Subsequently, samples were centrifuged at 12,000 x g for 20 min at 4 °C, and the clear upper layer was collected. To the obtained layer, 0.5 mL of isopropanol was added followed by a 10 min incubation at 4 °C. Subsequently, samples were centrifuged at 12,000 x g for 15 min at 4 °C, and the pellet was washed in 1 mL of cold 75% EtOH by centrifugation at 12,000 x g for 5 min at 4 °C, repeated twice. The collected pellet was air-dried for 10 min and resuspended in 30 μL of nuclease-free water. TURBO DNA-free Kit (Invitrogen) was used to eliminate the remaining genomic DNA from the isolated RNA samples. cDNA was synthesized from the isolated RNA with the High-Capacity cDNA Reverse Transcription Kit (Thermo Fisher Scientific). qPCR reactions were prepared using Maxima SYBR Green/ROX qPCR Master Mix (Thermo Fisher Scientific) with primer sets for Axin2 (F: 5’-CGACCCAGTCAATCCTTATCAC-3’, R: 5’-GGGACTCCATCTACGCTACTG-3’), Birc5 (F: 5’-CCAGGCATGAAGAGTCAGGG-3’, R: 5’-GGCTGCCTGCTTAGAGTTGA-3’), Ki67 (F: 5’-GAGGAGAAACGCCAACCAAGAG-3’, R: 5’-TTTGTCCTCGGTGGCGTTATCC-3’), Sprr1a (F: 5’-CAAGGCACCTGAGCCCTGCAA-3’, R: 5’-AGGCTCTGGTGCCTTAGGTTGG-3’), and Krt1 (F: 5’-GACTCGCTGAAGAGTGACCAGT-3’, R: 5’-GGTCACGAACTCATTCTCTGCG-3’) genes. Gene expression was normalized to Gapdh (F: 5’-TGGCAAAGTGGAGATTGTTGCC-3’, R: 5’-AAGATGGTGATGGGCTTCCCG-3’), and relative expression was calculated using the ΔΔCt method. Kruskal-Wallis test with Dunn’s multiple comparison test was used to find significant differences between the groups.

### Retroviral transduction mediated Runx3 overexpression

Retroviral particles encoding Runx3-IRES-EGFP or mCherry alone were produced as described previously ^12^. Briefly, Plat-E cells were seeded using high glucose DMEM (HyClone) supplemented with 10% fetal bovine serum in 6-well plates at a density of 5 × 10^5^ cells/well 1 d before transfection. Transfections were performed using 1.5 µg plasmid DNA from pRunx3-EGFP and 1 µg pCL-Eco with TransIT-LT1 (Mirus) in Opti-MEM I Reduced-Serum Medium (Gibco). Retroviral supernatant was harvested 48 h and 72 h after transfection. For transductions, negatively enriched naive CD8 T cells from spleen and lymph nodes were activated in 6-well plates coated with 100 µg/ml goat anti-hamster IgG (H+L; Thermo Fisher Scientific), 1 µg/ml anti-CD3 (145-2C11; eBioscience), and 1 µg/ml anti-CD28 (37.51; eBioscience). T cells were subsequently transduced by replacing media with retroviral supernatant supplemented with 50 µM β-mercaptoethanol (Gibco) and 8 µg/ml polybrene (Millipore) followed by a 1 h spinfection centrifugation at 2,000 rpm and 37°C. One day after transduction, Runx3 and empty vector transduced cells were mixed 1:1, and 100,000 total cells were co-cultured with organoids for 5-10 days to generate TRM.

### In vitro Peptide Restimulation Assay

After at least 7 days of VEO-CD8 T cell co-culture, the wells were treated with 0.2 μg/ml of cognate peptide (SIINFEKL for OT-I CD8 T cells, gp33 for P14 CD8 T cells, or SL8 for gBT-I CD8 T cells) for 4 hours in a restimulation media containing 10% FBS, 1% pennicililin/streptomycin, 1% L-glutamine, 1% non-essential amino acids, 1% sodium pyruvate, 0.25 ug/mL Amphotericin B, and 55μM beta-mercaptoethanol in RPMI, supplemented with Brefeldin-A. After 4 hours at 37°C, the cells were collected, washed, and stained for phenotype assessment using the BD cytofix-cytoperm system as per manufacturer’s instruction.

### Statistics

If the samples followed normal distribution, then parametric tests (unpaired two-tailed Student t-test for two groups and one-way ANOVA with Tukey multiple-comparison test for more than two groups) were used. Two-way ANOVA with Sidak multiple-comparison test was used if the effect of two independent variables were being considered among more than two sample groups. If the samples deviated from a Gaussian distribution, nonparametric tests (Mann-Whitney U test for two groups and Kruskal-Wallis with Dunn multiple-comparison test for more than two groups) were used, unless otherwise stated in the figure legends. For paired analyses not conforming to Gaussian distribution, the Wilcoxon matched-pair signed-rank test was used. Shapiro-Wilk normality test was used to determine whether samples adhered to Gaussian distribution or not. Variances between groups were compared using an F test and found to be equal. All statistical analysis was done in Prism (GraphPad Software). p values <0.05 were considered significant.

## Acknowledgments

Schematics were generated with Biorender.com. This work was supported by National Institutes of General Medical Sciences Grant 2P20GM109035, Rhode Island Foundation, Searle Scholar’s program (to L.K.B.). G.H. was supported by an American Association of Immunology Careers in Immunology Fellowship. F.J.M. was supported by Brown Respiratory Research Training Program NIH T32HL134625 and Molecular Biology, Cell Biology, and Biochemistry Graduate Program training grant 5T32GM136566. We would like to thank David Knipe (Harvard University) and Gregoire Lauvau (Albert Einstein College of Medicine) for providing the HSV-1gfp virus and gBT-I CD8 transgenic mice. We would also like to thank Peter Ruminski (Washington University) for synthesizing the CWHM-12 compound. We acknowledge the NIH tetramer core facility for providing all the tetramer reagents used in the study and Brown University flow cytometry core for facilitating the flow-based assays.

## Author contributions

Conceptualization: MRU, YL, LKB; Methodology and reagents: CM, SG, NZ, JM; Experimentation: MRU, YL, JRR, GH, MHH, FJM, LKB; Data analysis: MRU, YL, LKB; Writing: MRU, YL, LKB; Funding acquisition: LKB

